# KIF21B binds Myosin Va for Spine Entry and regulates Actin Dynamics to control Homeostatic Synaptic Downscaling

**DOI:** 10.1101/2022.05.04.490582

**Authors:** Kira V. Gromova, Edda Thies, Céline D. Dürst, Daniele Stajano, Michaela Schweizer, Marina Mikhaylova, Christine E. Gee, Matthias Kneussel

## Abstract

Homeostatic synaptic plasticity adjusts the strength of synapses to restrain neuronal activity within a physiological range. Postsynaptic GKAP controls the bidirectional synaptic scaling of AMPA receptors (AMPARs) however how chronic activity triggers postsynaptic protein remodeling to downscale synaptic transmission is barely understood. Here we report that the microtubule-dependent kinesin motor KIF21B interacts with GKAP and likewise enters dendritic spines in a myosin Va- and activity-dependent manner. We observed that under conditions of chronic activity KIF21B regulates actin dynamics in spines, triggers spine removal of GluA2-containing AMPA receptors, and mediates homeostatic synaptic downscaling of AMPA receptor-mediated mEPSC amplitudes. Our data highlight a myosin-kinesin interaction that enables the entry of the microtubule-dependent motor KIF21B into actin-rich spine compartments. A slow actin turnover rate might be beneficial for efficient protein removal from excitatory synapses, suggesting a functional role of KIF21B in a GKAP- and AMPA receptor-dependent mechanism, underlying homeostatic downscaling of neuronal firing.

## Introduction

Homeostatic synaptic plasticity is a feedback mechanism used by neurons and neuronal networks to balance excessive excitation or inhibition by adjusting synaptic strength^1^. It is suggested that long-term changes in neuronal firing rates induce changes in receptor trafficking to increase or decrease the number of glutamate receptors at synapses^2, 3^. Since homeostatic plasticity involves the global modification of synapses, it operates over longer timescales. At postsynaptic sites, several molecules including BDNF, CaMKII, Arc, Homer1a, GKAP or Shank are known to be involved in synaptic scaling^3^, however, the underlying signaling pathways and molecular mechanisms still need to be investigated^1^. Likewise, it is unknown whether and how the postsynaptic cytoskeleton rearranges following chronic activity and, which molecular motors drive for instance AMPAR removal during homeostatic synaptic scaling.

The guanylate kinase-associated protein (GKAP) is a postsynaptic scaffold component that links NMDA receptor/PSD-95 to Shank/Homer complexes^4^. Reduction of synaptic activity induces an accumulation of GKAP at excitatory spine synapses^3^, a process that requires interaction with myosin Va^5^. In contrast, overexcitation activates the calcium-calmodulin-dependent kinase CaMKII that promotes the synaptic removal of GKAP via degradation by the ubiquitin-proteasome system^3^. Trafficking mechanisms in dendritic spines that rearrange receptors, cell adhesion molecules and postsynaptic density proteins require a dynamic actin cytoskeleton, endocytic recycling mechanisms, and molecular motors to adjust synaptic strength^6, 7, 8^. In addition to actin rearrangement, microtubules transiently polymerize into dendritic spines, for instance, to deliver synaptotagmin IV via KIF1A^9, 10, 11^.

Motor proteins of the kinesin-4 family regulate microtubule dynamics and, in addition, may mediate processive transport^12, 13, 14, 15, 16^. Members include KIF4/Xklp1, which reduces the rate of microtubule growth and suppresses catastrophes^17^, the immotile motor KIF7 which also reduces microtubule growth but promotes catastrophes^18^, and the two large motors KIF21A and KIF21B^19^. Point mutations in the *Kif21a* gene cause the dominant eye movement syndrome Congenital Fibrosis of the Extraocular Muscles type 1 (CFEOM1)^20^, whereas KIF21B has been discussed as a risk factor in neurodevelopmental^12^ and neurodegenerative diseases^21, 22^. *Kif21b* haploinsufficiency in patients leads to impaired neuronal positioning and brain malformations^12^. In mice, the loss of KIF21B causes aberrant dendritic arborization of hippocampal CA1 pyramidal cells^23^, accompanied by learning and memory deficits such as in the initial encoding of spatial information, the memory of a tone-shock association^23^, and reduced cognitive flexibility^24^.

In physiology, KIF21B mediates different functions in individual cell types^25^, including neurons^12, 13, 15, 16, 23, 26^. It acts as a processive motor that, in response to neuronal activity contributes to the retrograde trafficking of brain-derived neurotrophic factor (BDNF)-TrkB complexes^16^ and participates in the cell surface delivery of GABAA receptors^14^. On the other hand, KIF21B regulates the dynamics of the microtubule cytoskeleton by accumulating at microtubule plus ends, thereby pausing microtubule growth^13, 26^. Consequently, *Kif21b*-deficient neurons are characterized by longer microtubules, than wild-type control cells^23^.

At excitatory synapses, KIF21B mediates the translocation of a Rac1 guanine nucleotide exchange factor (ELMO1) from dendritic spines to terminate Rac1 activity, a process underlying the expression of long-term depression (LTD)^24^. However, although KIF21B regulates proteins at dendritic spines, it has remained unclear whether the motor itself can enter spine protrusions and whether the actin cytoskeleton might be involved in KIF21B-dependent processes.

Here, we report that the kinesin KIF21B directly interacts with myosin Va and enters the actin-rich dendritic spine compartment in a myosin Va-dependent manner. Following chronic stimulation, we show that KIF21B functionally associates with GKAP and is a prerequisite for the dynamic rearrangement of actin filaments upon the induction of homeostatic synaptic downscaling.

## Results

### The kinesin KIF21B enters dendritic spine protrusions

The kinesin motor and microtubule pausing factor KIF21B, is a critical player in regulating spine morphology, neuronal function, and behavior ^16, 23, 24, 27^. Whether the kinesin is restricted to neuronal dendrites or enters dendritic spine synapses has remained unknown. Using KIF21B-specific antibodies, verified by *Kif21b* knockout brain tissue, we identified the endogenous kinesin at the tips of individual F-actin-rich spine protrusions colocalized with the postsynaptic density protein PSD-95 in cultured hippocampal neurons (Figure 1A-C). About 55% of the F-actin-labeled spines were double-positive for KIF21B and PSD-95 (Figure 1D). Biochemical fractionation to enrich postsynaptic densities (PSDs), known to contain PSD-95 and GKAP^4, 28^ but just small amounts of the presynaptic marker synaptophysin, further identified KIF21B in PSD fractions (Figure 1E). We therefore combined diaminobenzidine (DAB)-staining with immuno-electron microscopy to assess the subcellular distribution of KIF21B. Consistent with KIF21B regulating microtubule growth and microtubule-mediated transport^13, 16, 23, 24, 27^, the DAB-labeled motor protein decorated dendritic microtubules in wildtype, but not in *Kif21b* knockout tissue (Figure 1F, left and box 1). In addition, KIF21B was detected at the PSD of individual excitatory spine synapses in the hippocampus (Figure 1F, right and box 2). We therefore conclude that the kinesin is not restricted to dendritic microtubules, but can be located postsynaptically in an F-actin-rich compartment.

**Figure 1.**
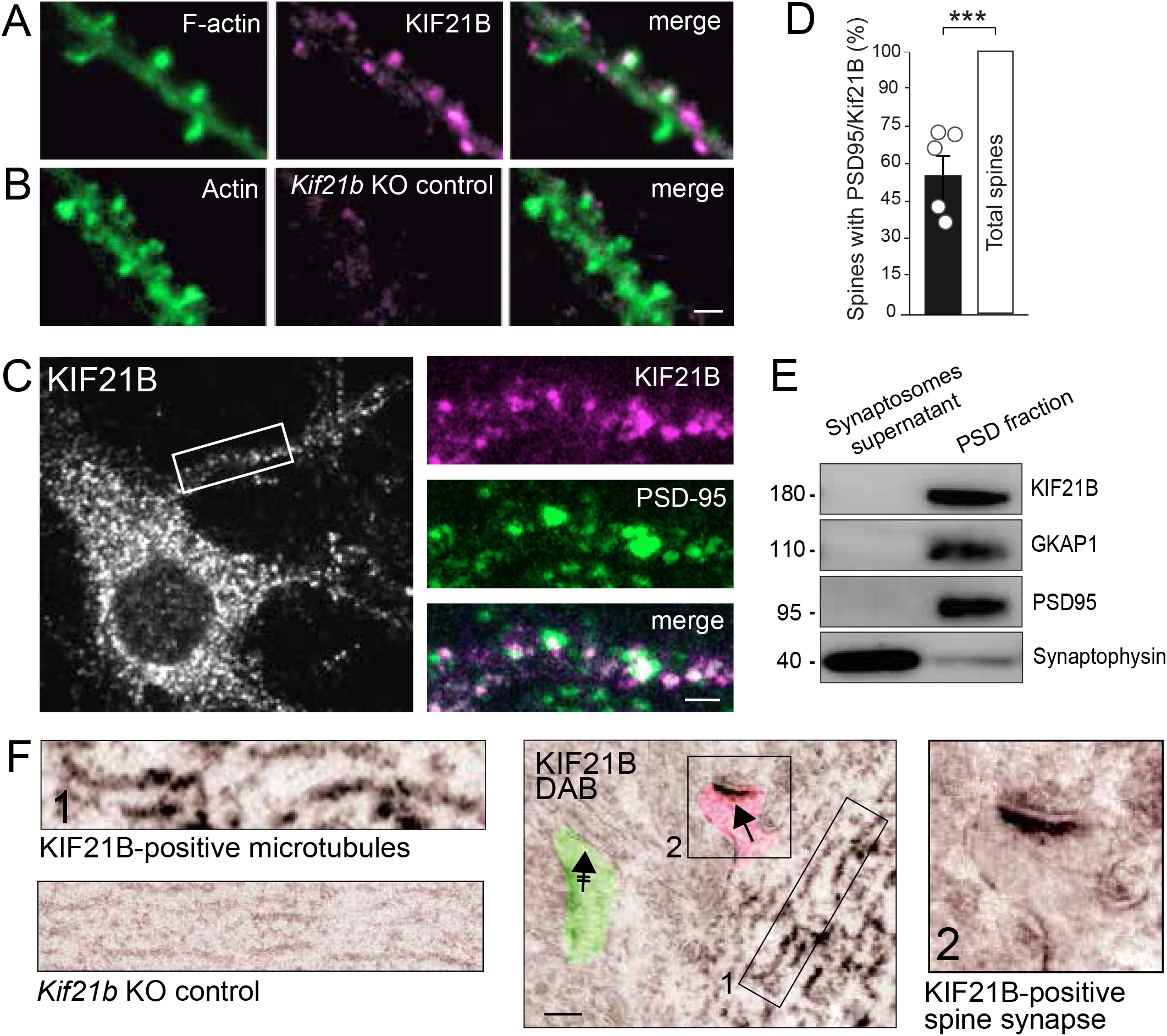
Kif21B is located at postsynaptic sites of dendritic spines. (**A, B**) DIV14 cultured hippocampal neurons from KIF21B^+/+^ (**A**) and KIF21B^−/−^(**B**) mice, co-labeled with F-actin-(green) or Kif21B-(magenta) specific antibodies. (**C**) Wildtype DIV14 cultured hippocampal neurons co-labeled with Kif21B-(magenta) or PSD-95-(green) specific antibodies. (**D**) Quantification of C (n=1,400 spines; N=5 exp.). Data are expressed as a percentage of control (total spine number) ± SEM. Statistical analyses: one-sample t-test; t=10,7 df=6, p=0,0001. (**E**) Western blot analysis of Kif21B in postsynaptic density fractions from mouse hippocampal tissue. Presynaptic synaptophysin is enriched in the supernatant (synaptosomes). KIF21B, GKAP1, and PSD-95 are detected in postsynaptic fractions (N=3). (**F**) Immunoelectron microscopy with diaminobenzidine (DAB) using hippocampal slices. Boxed region 1 and left: KIF21B decorates microtubules in KIF21B^+/+^ (upper) but not KIF21B^−/−^ neurons (KO control, lower). Boxed region 2 and right: KIF21B is detected at the PSD of individual spines. KIF21B-positive spine (pink, arrow), KIF21B-negative spine (green, crossed arrow). Scale bars: 3 µm (**A-C**), 500 nm (**F**).

### Myosin Va and neuronal activity changes regulate the localization of KIF21B in spines

Microtubules occasionally polymerize into dendritic spines^9, 11^ and invading microtubules mediate KIF1A-mediated transport into spine compartments^10^. However, the number of spines containing microtubules is much lower^9, 11^, as compared to the number of spines containing KIF21B. We therefore asked whether the kinesin might enter spine protrusions independent from microtubules. Our previous proteomic screen had identified the unconventional myosin Va (MyoVa), as a potential KIF21B binding partner^27^, a highly abundant actin-based motor mediating AMPAR^29^ or GKAP^5^ trafficking. Remarkably, co-immunoprecipitation (co-IP) from hippocampal lysate confirmed a specific interaction of the kinesin KIF21B with myosin Va, but not with myosin Vb. MyoVa-specific antibodies precipitated endogenous MyoVa and led to co-IP of endogenous KIF21B (Figure 2A). Vice versa, KIF21B-specific antibodies precipitated the endogenous kinesin and led to co-IP of endogenous myoVa (Figure 2B). Interestingly, this interaction was only identified in the presence of phosphatase inhibitor, suggesting a phospho-dependent regulation. To assess whether the kinesin directly binds the myosin, we employed the heterologous DupLex-A yeast two-hybrid assay. Using different deletion mutants of *Kif21b* and *MyoVa*, we identified the N-terminal region of KIF21B, containing the motor domain and parts of the stalk domain, to mediate interaction with the C-terminal region of MyoVa, harboring its coiled-coil and tail domains (Figure 2C, D). Since the tail domain of MyoVa is typically involved in cargo binding^6^, we asked whether chemical inhibition of MyoVa-mediated transport^30^ altered the localization of KIF21B in F-actin/PSD-95 double positive spines. While inhibition of MyoVa over 1 h significantly reduced the number of spines containing KIF21B (Figure 2E), inhibition of myosin Vb or the use of the microtubule polymerization inhibitor nocodazole had no effect (Figure 2F), suggesting that KIF21B enters dendritic spines influenced by MyoVa but independent of MyoVb or microtubules.

**Figure 2.**
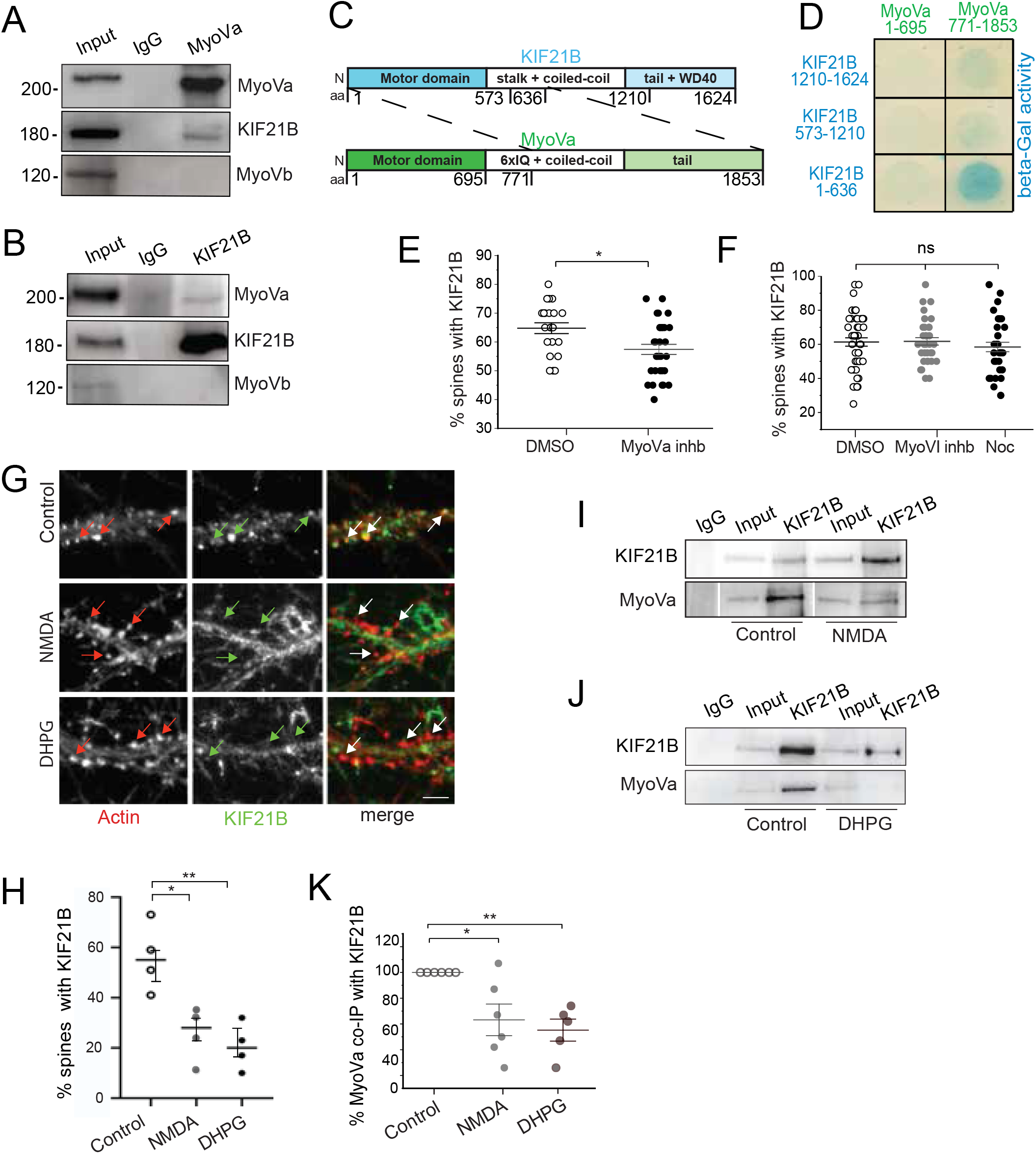
The localization of KIF21B in spines is myosin Va and synaptic activity-dependent. (**A**) Co-IP with myosin Va (MyoVa)-specific antibodies using adult mouse hippocampal lysate. Detection of MyoVa, KIF21B, and Myosin Vb (MyoVb). IgG: control (N=4). (**B**) Co-IP with Kif21B-specific antibodies using adult mouse hippocampal lysate. Detection of MyoVa, KIF21B, and MyoVb. IgG: control (N=4). (**C**) Schematic domain representation of fragments used in **D**. (**D**) Yeast-two-hybrid screen of KIF21B (prey) with MyoVa (bait) (N=2). (**E**) Quantification of KIF21B-positive spines co-labeled with F-actin and PSD95 following treatment with DMSO (control, n=22 cells, n=440 spines) or MyoVa inhibitor (n=35 cells, n=700 spines); N=3 experiments, Student’s t-test t=2,163 df=54, p=0,035. (**F**) Quantification of KIF21B-positive spines co-labeled with F-actin and PSD95 following treatment with DMSO (control, n=49 cells, n=980 spines), MyoVI inhibitor (n=31 cells, n=620 spines) or nocodazole (Noc, n=36 cells, n=720 spines), N=3 experiments, One-Way ANOVA Post-hoc Tukey, F=1,098 df=111. p= 0,97 (DMSO/MyoVI), p=0,68 (DMSO/Noc), p=0,62 (MyoVI/Noc). (**G**) Hippocampal neurons co-labeled with KIF21B-(green) and F-actin-(red) specific antibodies, following treatment with NMDA or DHPG. (**H**) Quantification of KIF21B-positive spines from G. Control: n=34 cells, n=505 spines. NMDA: n=30 cells n=437 spines. DHPG n=26 cells, n=354 spines. N=4 experiments, each. One-Way ANOVA, Post-hoc Tukey, F=11,5 df=11, p=0,01 (Control/NMDA), p=0,004 (Control/DHPG). (**I**) Co-IP with KIF21B-specific antibodies using acute hippocampal slices treated with NMDA. Detection of myosin Va (MyoVa). N=5. (**J**) Co-IP with KIF21B -specific antibodies using acute hippocampal slices treated with DHPG. Detection of myosin Va (MyoVa). N=5. (**K**) Quantification of **I** and **J** (N=5). One-Way ANOVA, Post-hoc Dunnett F=11,1 df=14, p=0,01 (Control/NMDA), p=0,0057 (Control/DHPG). Data expressed as a percentage of control ± SEM.

Since KIF21B has been implicated in Rac1 regulation underlying LTD expression^24^, we further asked whether neuronal activity changes could in general affect the spine localization of the kinesin. To this end, we applied two independent chemical protocols to induce LTD (cLTD). Treatment of cultured hippocampal neurons with either 40µM NMDA for 10 minutes^31^ or 50µM DHPG for 30 minutes^32^ significantly reduced the number of KIF21B-positive spines, as compared to control conditions (Figure 2G, H). Based on our hypothesis, we further treated acute hippocampal slices with these drugs and performed co-IP experiments, using KIF21B-specific antibodies. Accordingly, both cLTD protocols significantly reduced the amount of myosin Va that coprecipitated with the kinesin (Figure 2I-K), indicating that the interaction of both motor proteins and consequent the delivery of KIF21B into dendritic spines are activity-dependent processes.

### Knockout of the *Kif21b* gene alters F-actin dynamics in dendritic spines

As in former studies^23, 24^, the size of Dil-labeled dendritic spines in cultured hippocampal neurons (Figure 3A, B) or in area CA1 of hippocampal slices (Figure 3C, D) was significantly increased in *Kif21b* knockouts under basal conditions. Since changes in spine morphology are often attributed to the actin cytoskeleton^33^, we co-expressed KIF21B-GFP and mRFP-actin in COS-7 fibroblasts. Under control conditions, the kinesin appeared with a prominent localization at the periphery of these cells (Figure 3E and Supplemental Movie 1). Remarkably, in cells treated with the actin polymerization inhibitor cytochalasin D (CytoD), punctate mRFP-actin signals revealed a strong colocalization with KIF21B-GFP (Figure 3F and Supplemental Movie 2) that was not apparent in control cells (Figure 3E). As CytoD increases the number of F-actin ends^34^, KIF21B might preferentially interact with the ends of actin filaments.

**Figure 3.**
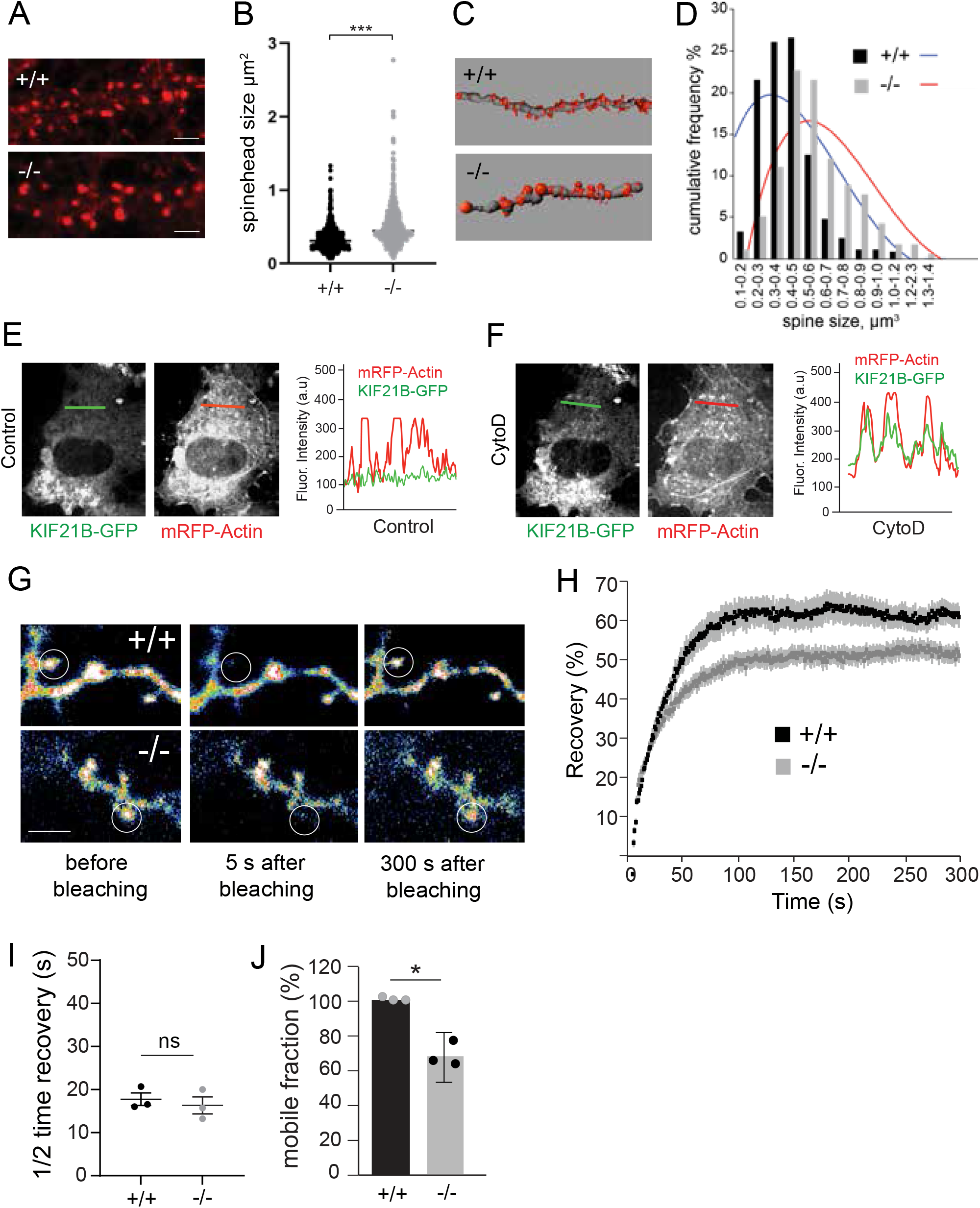
Kif21B depletion affects spines size and actin stability in spines. (**A**) Dendritic protrusions from Dil-labeled KIF21B^+/+^ and KIF21B^−/−^ cultured hippocampal neurons at DIV14. (**B**) Quantification of spine head size per length of dendrites (n= 600 spines, N=3 experiments). Mann-Whitney U Test, p=0,0001. (**C**) Dendritic protrusions from Dil-labeled KIF21B^+/+^ and KIF21B^−/−^ hippocampal slice cultures from CA1. 3D reconstruction (KIF21B^+/+^: n=73, KIF21B^−/−^: n=181 spines, N=3 experiments). (**D**) Cumulative frequency (%) of spine head size (µm^3^) per dendrite length from C. (**E, F**) Localization of mRFP-Actin and KIF21B-GFP in COS7 cells treated with cytochalasin D (CytoD) for 1h, n=10 cells, each. Right: Line scans depict overlapping fluorescent signal intensities (a.u.) over the lines in E (control) and F (CytoD). (**G**) Spine heads one day after transfection with GFP-actin using KIF21B^+/+^ or KIF21B^−/−^ cultured hippocampal neurons at DIV13. FRAP imaging of individual spines (circles) before and after bleaching is indicated. (**H**) FRAP analysis of GFP-actin fluorescence recovery. Recovery in KIF21B^−/−^ spines (grey) does not reach the same level as in KIF21B^+/+^ (black). Equal imaging and photobleaching conditions were used. (**I**) Average GFP-actin half-time recovery (s) of the dynamic F-actin pool is comparable in spines derived from KIF21B^+/+^ and KIF21B^−/−^ neurons. Mann-Whitney U Test, p=0,594. (**J**) The mobile F-actin fraction in FRAP recovery curves is significantly decreased in KIF21B^−/−^ spines, indicating increased actin stability. KIF21B^+/+^ (n=99 spines), KIF21B^−/−^ (n=135 spines); N=3 experiments. One sample t-test, t=5, df=2, p=0,0375. Scale bar = 5s µm. Data are expressed ± SEM.

To address the dynamics and turnover of the synaptic actin cytoskeleton, we employed fluorescence recovery after photobleaching (FRAP), using DIV13 cultured hippocampal neurons from *Kif21b*^+/+^ and *Kif21b*^−/−^ mice expressing GFP-actin. Selective photobleaching of single spines in both control and knockout neurons rapidly decreased the fluorescence of GFP-actin and led to a fast recovery of the signals (Figure 3G-I). However, the GFP-actin pool in *Kif21b* knockout neurons did not recover to the same level (Figure 3G, lower images, H), suggesting that in the absence of *Kif21b* gene expression, dendritic spines contain more stable actin and a significantly reduced mobile fraction (Figure 3J). We therefore conclude that the kinesin KIF21B participates in the regulation of actin dynamics at spine synapses.

### KIF21B associates with GKAP and participates in the regulation of homeostatic synaptic downscaling

GKAP is a postsynaptic protein connecting actin filaments with the PSD^4^. Since GKAP also interacts with a light chain of MyoVa^5^, we asked whether KIF21B and GKAP might be functionally associated. Co-IP experiments from mouse brain lysate using GKAP-specific antibodies precipitated GKAP and led to co-precipitation of KIF21B, Shank and Homer, but not the related kinesin motor KIF21A (Figure 4A and Figure S1A). Reciprocal co-IPs using KIF21B-specific antibodies confirmed the GKAP-KIF2B interaction (Figure S1B). Accordingly, endogenous GKAP and KIF21B were frequently colocalized in actin-positive dendritic protrusions (Figure 4B), with about 30% of all spines depicting colocalization of both proteins. GKAP is a prominent molecular player regulating homeostatic synaptic scaling and is displaced from synaptic spines following chronic stimulation^3^. We therefore employed established homeostatic synaptic plasticity (HSP) protocols^35, 36^, to ask whether KIF21B might also participate in homeostatic regulation. Following chronic stimulation with the GABA_A_ receptor antagonist bicuculline (BIC) or activity blockade with the sodium channel blocker tetrodotoxin (TTX) over 48 h, we analyzed the fluorescence intensity of GluA2-type AMPAR immunoreactivity at the cell surface of spines, applying a receptors surface staining protocol in the presence (+/+) or absence (-/-) of KIF21B (Figure 4C, D). Quantification of wildtype (+/+) neurons, confirmed published data, which show that BIC treatment significantly reduces surface AMPARs over 48h, whereas TTX treatment significantly increases them^35, 36^ (Figure 4C, E). In contrast, in *Kif21b* knockout (-/-) neurons, chronic BIC treatment led to opposite results, increasing spine surface AMPARs, whereas chronic TTX had no significant effect (Figure 4D, F).

**Figure 4.**
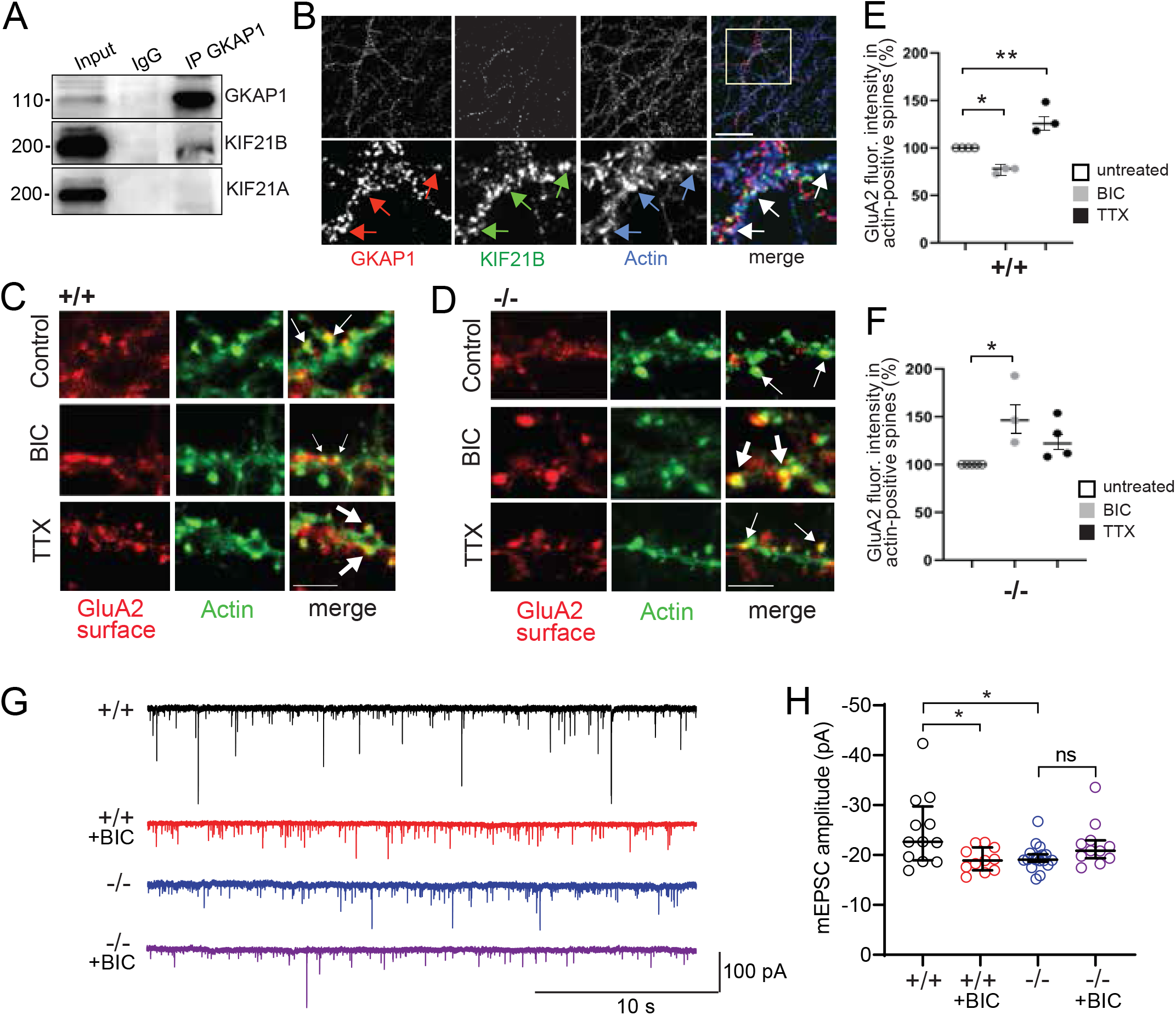
KIF21B associates with GKAP and regulates homeostatic synaptic downscaling. (**A**) Co-IP with GKAP-specific antibodies using adult mouse brain lysate. Detection of GKAP1, KIF21B, and KIF21A (negative control), N=3. (**B**) Hippocampal neurons at DIV14 co-labeled with GKAP-(red), KIF21B-(green) and F-actin-(blue) specific antibodies. Arrows depict colocalization, n=45 cells, N=3 experiments. (**C, D**) Hippocampal neurons at DIV14 co-labeled with a surface staining protocol using GluA2-(red) and F-Actin-(green) specific antibodies, following treatment with BIC or TTX over 48 hours. (**C**) KIF21B^+/+^ wildtype neurons. (**D**) KIF21B^−/−^ knockout neurons. Arrows depict colocalization. The thickness of arrows corresponds to values in E and F. (**E, F**) Quantification of cell surface GluA2 fluorescence intensity in spines from C and D (n=30 cells; N≥3 experiments, each. One-way ANOVA, Post-hoc Dunnett; for KIF21B^+/+^ F=30 df=9 and KIF21B^−/−^ F=3,7 df=11, (E) p=0,016 (Untreated/BIC), p=0,004 (Untreated/TTX). (F) p=0,01 (Untreated**/**BIC). (**G**) Representative mEPSC recordings using KIF21B^+/+^ or KIF21B^−/−^hippocampal slices treated with BIC over 48 hours. (**H**) Quantification of mEPSC amplitudes shown in G, (pA), ANOVA, Post-hoc Tukey. p=0,0365 (Control (+/+)/BIC (+/+)), p=0,0406 Control (+/+)**/**Control (-/-), p=0,2466 (Control (-/-)/BIC (-/-)). n=12 slices (Control+/+), n=11 slices (BIC+/+), n=18 slices (Control-/-), n=11 slices (BIC-/-). Data are expressed ± SEM.

We then recorded AMPAR-mediated miniature excitatory postsynaptic currents (mEPSCs) from CA3 neurons in DIV 23-24 hippocampal slice cultures. In this assay, BIC is known to increase network activity and, upon chronic application, to induce homeostatic downscaling of AMPA mEPSCs^35, 36, 37^. Accordingly, chronic BIC application significantly decreased mEPSC amplitudes in wildtype slices (Figure 4G, H, black vs. red), without changing mEPSC frequency (Figure S1C). The amplitude of mEPSCs from untreated *Kif21b* knockout neurons were significantly smaller than from control slices with no change in frequency (Figure 4G, H, blue vs. black and Figure S1C). In CA3 neurons from slices deficient in KIF21B, chronic BIC application failed to induce any reduction of mEPSC amplitude or frequency (Figure 4G, H, blue vs. purple and Figure S1C). Together, these data suggest that KIF21B, similar to its binding partner GKAP, is a critical determinant in the homeostatic regulation of synaptic AMPAR levels and participates in homeostatic synaptic downscaling.

### The regulation of homeostatic synaptic downscaling though GKAP requires KIF21B

In addition to AMPARs, homeostatic downscaling decreases the levels of other postsynaptic proteins^37^. In part, these decreases are mediated through ubiquitination and the subsequent proteasomal degradation of GKAP^3^. In order to check whether GKAP leaves dendritic spines under the conditions of our study, we quantified fluorescence intensity of immunostained endogenous GKAP and F-actin, following 48h of BIC. Consistent with the literature^3^, GKAP significantly decreased at actin-positive spine protrusions, as compared to control conditions (Figure 5A, B). Notably, the GKAP binding partner KIF21B also was reduced significantly at spines after chronic BIC (Figure 5C, D), supporting our hypothesis that the kinesin might participate in mechanisms of homeostatic synaptic downscaling. To assess whether the synaptic removal of GKAP might be KIF21B-dependent, we performed this assay in the absence of *Kif21b* gene expression. Whereas chronic BIC treatment significantly reduced GKAP from actin-positive protrusions in wildtype neurons, expressing KIF21B (Figure 5E, F), the effect was abolished in *Kif21b* knockout neurons (Figure 5F, G). Thus, the functional role of GKAP in the regulation of synaptic downscaling^3^ requires the kinesin KIF21B.

**Figure 5.**
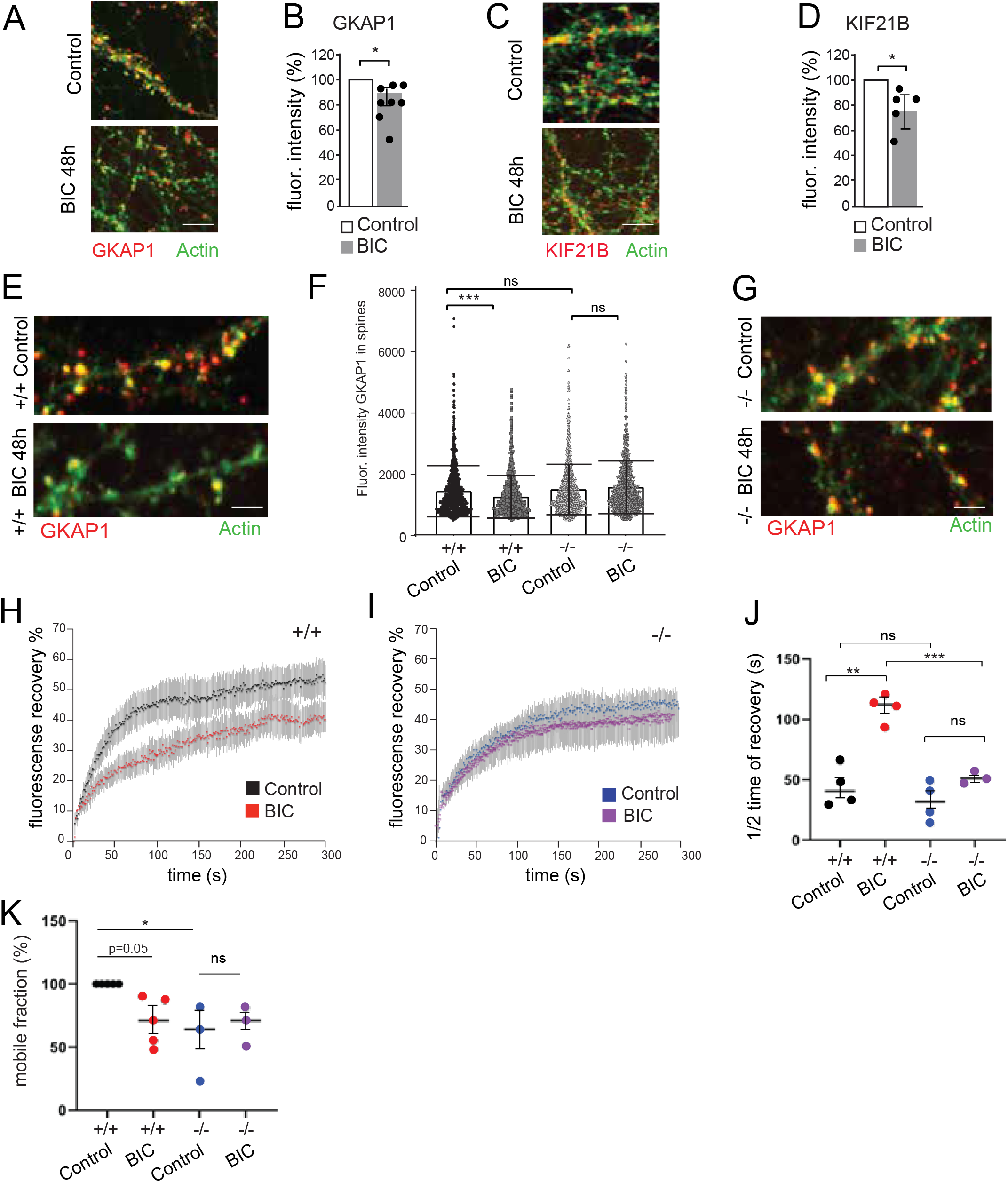
KIF21B regulates GKAP and actin dynamics in spines following the induction of homeostatic synaptic plasticity. (**A**) Hippocampal neurons at DIV14 co-labeled with GKAP-(red) and F-Actin-(green) specific antibodies, following treatment with DMSO (control) or BIC over 48 hours. (**B**) Quantification of fluorescent intensity (%) of GKAP1 in F-actin-positive spines in A (n= 120 cells, n=8 experiments). One sample T-test, t=3,1 df=7, p=0,0169. Data are expressed ± SEM. (**C**) Hippocampal neurons at DIV14 co-labeled with KIF21B-(red) and F-Actin-(green) specific antibodies, following treatment with DMSO (control) or BIC over 48 hours. (**D**) Quantification of fluorescent intensity (%) of KIF21B in F-actin-positive spines in C (n=75 cells, n=5 experiments). One sample T-test, t=2.86 df=4, p=0,045. Data are expressed ± SEM. (**E, G**) Hippocampal neurons at DIV14 derived from KIF21B^+/+^ (E) or KIF21B^−/−^ (G), co-labeled with GKAP-(red) and F-Actin- (green) specific antibodies. Neurons were treated with either DMSO (control) or BIC over 48. (**F**) Quantification of GKAP fluorescence intensity from E and G (n=1,110 spines for each condition, N=4 experiments). One-way ANOVA, Post-hoc Bonferroni; F=23,8 df=4406, p=0,0001 (Control (+/+)**/**BIC (+/+)), p=0,534 (Control (+/+)**/** Control (-/-)), p=0,1966 (Control (-/-)**/**BIC (-/-)). Data are expressed as individual data points. (**H**) FRAP analysis of GFP-actin recovery using wildtype (KIF21B^+/+^) hippocampal neurons at DIV 13 treated with DMSO (control) or BIC over 48h. (**I**) FRAP analysis of GFP-actin recovery using knockout (KIF21B^-/-^) hippocampal neurons at DIV 13 treated with DMSO (control) or BIC over 48h. (**J**) The average GFP-actin half-time recovery (s) of the dynamic F-actin pool in spines is significantly increased in KIF21B^+/+^ neurons, following treatment with BIC, but remains unchanged in KIF21B^−/−^ neurons. One-way ANOVA, Post-hoc Tukey F=25,9 df =14, p=0,001 (Control (+/+)**/**BIC (+/+)), p=0,56 (Control (+/+)**/** Control (-/-)), p=0,0008 (BIC (+/+)**/** BIC (-/-)), p=0,26 (Control (-/-)**/** BIC (-/-)). (**K**) The mobile actin fraction measured from FRAP curves of individual spines is by trend slightly decreased in KIF21B^+/+^ spines after treatment with BIC but remains unchanged for KIF21B^−/−^ spines. One-way ANOVA, p=0,05 (Control (+/+)**/**BIC (+/+)), p=0,0128 (Control (+/+)**/** Control (-/-)), p=0,844 (Control (-/-)**/** BIC (-/-)). (**J, K**) KIF21B^+/+^ DMSO (n=55 spines) N≥4 experiments, KIF21B^+/+^ BIC (n=69 spines) N≥4 experiments, KIF21B^−/−^ DMSO (n=80 spines) N≥3 experiments, KIF21B^−/−^ BIC (n=59 spines) N≥3 experiments. One-way ANOVA, Post-hoc Tukey, F=3,1 df=16. Scale bars:10 µm (A, C), 5 µm (E, G).

Finally, we aimed to assess whether the induction of synaptic downscaling altered the dynamics of the actin cytoskeleton. To this end, we applied FRAP experiments with GFP-actin following chronic BIC treatment over 48h. In wildtype (+/+) neurons, F-actin turnover rates were significantly slower in the presence of BIC (Figure 5H and Figure S1D) characterized by a significantly longer half-time recovery (Figure 5J) and a tendency pointing to a reduced mobile fraction (Figure 5K). Remarkably, in the absence of *Kif21b* gene expression (knockout -/-neurons), this change in actin dynamics was not detectable (Figure 5I-K and Figure S1D), indicating that the kinesin is a prerequisite for the dynamic rearrangement of actin filaments upon the induction of synaptic downscaling.

## Discussion

This study reports that the kinesin motor protein KIF21B binds to the postsynaptic scaffold protein GKAP (Figures 4 and S1), a critical regulator of homeostatic synaptic scaling^3^. Like GKAP^3, 5^, KIF21B enters dendritic spines in a myosin Va-dependent manner (Figures 1 and 2). Under conditions of chronic activity that induce homeostatic synaptic downscaling (BIC over 48h), the kinesin stabilizes the actin cytoskeleton and reduces polymerization of F-actin in spines (Figure 5 and S1). This process seems to be critical for synaptic protein removal following the induction of homeostatic plasticity, since KIF21B is required for the removal of GKAP and GluA2-containing AMPARs from spines and for homeostatic synaptic downscaling of AMPAR-mediated mEPSC amplitudes (Figure 4 and 5). Our data connect a kinesin motor and microtubule pausing factor with synaptic homeostasis. They further demonstrate that a kinesin cooperates with a myosin and participates in the dynamic regulation of the actin cytoskeleton in dendritic spines.

So far, only a limited number of molecules and mechanisms have been shown to regulate homeostatic synaptic plasticity (HSP). At glutamatergic synapses, HSP regulation includes the delivery and removal of AMPA receptors to the postsynaptic plasma membrane^38, 39^. In addition to AMPA receptor diffusion within the plane of the plasma membrane^40, 41^, AMPA receptor turnover requires endocytosis, motor proteins and a highly regulated actin cytoskeleton in spines^29, 42, 43^. Whereas, AMPA receptors undergo endocytic recycling and eventually reinsert into the cell surface membrane^44, 45^, other postsynaptic factors leave the synapse via ubiquitination and subsequent proteasomal degradation. For instance, the postsynaptic density protein GKAP undergoes activity-dependent ubiquitination, which can be induced through the phosphorylation of CaMKII^3^. Notably, previous studies revealed GKAP as well as the kinesin KIF21B as interactors of the ubiquitin ligase TRIM3^46, 47^, suggesting that the ubiquitin-proteasome pathway might be involved in the processes described in this study.

Different studies reported physical binding^48, 49, 50^ or functional interactions^51^ between kinesin and myosin motor proteins. Dendritic spines mainly contain actin filaments^8^ and myosin motors^6, 29, 42^, however depending on neuronal activity, some dynamic microtubules occasionally polymerize into dendritic spines^9, 11^ and regulate cargo transport through kinesin motors, such as the KIF1A-dependent delivery of synaptotagmin IV^10^. Also, KIF5 participates in the synaptic removal of an AMPAR-protrudin complex from spines, following LTD expression^52^. In contrast to microtubule-dependent spine entry of kinesins, our study reports a novel association of KIF21B with myosin Va that points to KIF21B spine entry in a myosin Va-dependent manner. Since the number of KIF21B-positive spines exceeds the number of microtubule-positive spines by an order of magnitude^9, 11^, KIF21B might undergo piggyback transport by myosin Va. Consistent with this view, a chemical inhibitor of myosin Va interferes with the spine localization of KIF21B (Figure 2E).

Our finding that the kinesin KIF21B regulates the fluorescence recovery of actin after photobleaching was unexpected and might be an indirect effect involving other proteins. However, a kinesin from plants was shown to bind to actin filaments^53^ and the kinesin-like proteins KAC1/2 regulate actin dynamics underlying chloroplast light avoidance in plants^54^. In dendritic spines, actin filaments connect to the PSD-95-positive postsynaptic density through SynGAP, GKAP, Shank and cortactin, with GKAP being in an intermediate position^4^. Following HSP-triggered removal of GKAP from postsynaptic sites, the physical connection between actin filaments and the postsynaptic density might be weakened. Previous studies reported that the induction of synaptic plasticity causes rearrangement of actin filaments and the regulation of actin dynamics in spines^8, 55^. Our finding that KIF21B regulates spine actin stability following HSP induction could suggest that stable actin filaments and slower actin dynamics are a prerequisite for the efficient removal of spine proteins under HSP conditions.

KIF21B is involved in the regulation of critical mechanisms underlying synaptic plasticity, learning and memory^16, 23^. Previous data further reported that LTD-causing stimuli induce the dynamic association of KIF21B with the Rac1GEF subunit engulfment and cell motility protein 1 (ELMO1), leading to ELMO1 translocation out of dendritic spines and its sequestration into endosomes. Despite these functional knock-down and knockout studies, KIF21B was so far not detected at the PSD of spine protrusions. The present study is consistent with a synaptic role of the kinesin and has revealed that KIF21B not only regulates Hebbian, but also homeostatic processes at glutamatergic synapses. Future studies will have to clarify whether these different regulatory modes underlie overlapping molecular pathways and whether they require KIF21B as cargo transporter or pausing factor of microtubule growth.

Finally, the kinesin KIF21B is highly upregulated in neurodegenerative disease (NDD) including multiple sclerosis (MS) and Alzheimer’s disease (AD)^22^. Under these conditions, a high prevalence of epileptiform activity emerges as a common pathophysiological hallmark^56^. Upregulation of *Kif21b* gene expression was most prominent in younger AD patients up to 62 years of age, suggesting that KIF21B might be beneficial at early stages when neurodegeneration is still limited. However so far, no functional role of KIF21B could be related to NDDs. Our observation that KIF21B participates in the homeostatic synaptic downscaling of chronically elevated neuronal activity, could be a potential explanation for the upregulation of KIF21B in NDD. Whether this effect is an attempt of neurons to compensate for hyperexcitability in disease, requires further investigation. Nevertheless, our study provides a starting point to address these questions.

In summary, the presently available data about KIF21B in neurons reveal that the motor acts at a critical position underlying long-term depression^24^ and homeostatic synaptic downscaling at synaptic sites. To connect these physiological functions with synaptopathy will become a future challenge.

## Methods

### Antibodies and DNA Constructs

The following antibodies were obtained from commercial sources. Rabbit anti-KIF21B (1:1,000; Sigma-Aldrich, Taufkirchen, Germany), Mouse anti-Homer1 (1:1,000; Synaptic Systems, Göttingen, Germany), mouse anti-GluA2 extracellular (1:1,000; Millipore, Darmstadt, Germany), rabbit anti-MyoVa (1:1,000; Sigma-Aldrich, Taufkirchen, Germany), rabbit anti-MyoVI (1:1,000; Sigma-Aldrich, Taufkirchen, Germany), mouse anti-PSD95 (1:100; Thermo Fisher Scientific, Dreieich, Germany), goat anti-Shank1-3 (1:1,000; Abcam, Cambridge, UK), guinea pig anti-Synaptophysin (1:1,000; Synaptic Systems, Göttingen, Germany), rabbit anti-KIF21A (1:1,000; Dianova, Hamburg, Germany). Cy2, Cy3, Cy5, DyeLight conjugated donkey anti-rabbit, anti-mouse or anti-goat (1:1,000; Dianova, Hamburg, Germany), peroxidase-conjugated donkey anti-rabbit, anti-mouse, anti-goat (1:15,000; Dianova, Hamburg, Germany). The following constructs have been previously described: KIF21B-EGFP^13^; mRFP/EGFP-actin (Addgene, Cambridge, MA).

### Animals

Mice were housed and bred at the animal facility of the University Medical Center Hamburg-Eppendorf. All procedures were performed in compliance with German law and according to the guidelines of Directive 2010/63/EU. Protocols were approved by the “Behörde für Justiz und Verbraucherschutz, Lebensmittel und Veterinärwesen Hamburg”.

### Immunoprecipitation

All steps were carried out at 4°C. 30 µl “Dynabeads Protein G” (Life Technologies, Darmstadt, Germany) were washed in PBS and incubated with 2-5 µg of specific antibody or control IgG for 30-60 min. After washing in PBS and IM-Ac-buffer (20 mM HEPES, 100 mM K-Acetate, 40 mM KCl, 5 mM EGTA, 5 mM MgCl_2_, 1% Triton-X-100, 1x Complete Protease Inhibitor Cocktail (Roche, Mannheim, Germany), 1mM PMSF, 5 mM DTT and 2 mM ATP; pH 7.2) and 1x Phosphatase Inhibitor Cocktail (Roche, Mannheim, Germany), antibody-coupled beads were incubated for 2-4h or other night with mouse brain lysate. Beads were then extensively washed with IM-Ac-buffer, boiled in SDS sample buffer and analysed by western blotting. Brain lysates were obtained by differential centrifugation from whole mouse brains at postnatal day P23 or from isolated hippocampus P365, as described ^57^.

### Western blot analysis

Samples were incubated for 5 min at 95°C in SDS loading buffer and subjected to sodium dodecyl sulphate-polyacrylamide gel electrophoresis (SDS-PAGE). Proteins were transferred to polyvinylidene difluoride (PVDF) membranes using a semi-dry blotting system. Membranes were blocked in 3% BSA (bovine serum albumin) prior to overnight incubation with primary antibodies at 4°C. Membranes were then washed and incubated with secondary antibodies coupled to horseradish peroxidase (HRP). Immunoreactive bands were visualized using the chemiluminescence detection system (INTAS Chemo Cam 3.2, Göttingen, Germany). Optical densities of respective bands were analysed using the Image J software (NIH, Bethesda, MD).

### Primary hippocampal cultures, transfection, immunochemistry

Primary hippocampal neurons were prepared from embryonic day 16 (E16). Briefly, 12 mm coverslips were coated with poly-L-lysine (5 µg/ml in PBS). 60,000 cells were seeded per coverslip in Lonza PNGM medium (Thermo Fisher Scientific, Dreieich, Germany). Neurons were cultured for 14-24 days *in vitro* (DIV) and were transfected using a calcium phosphate precipitation protocol. Briefly, per 22 mm coverslip, 2 μg of DNA (250 mM CaCl_2_ in 25 μl) was mixed with 25 μl of 2x HBS (42 mM HEPES, 10 mM KCl, 12 mM dextrose, 274 mM NaCl, 1.5 mM Na_2_PO_4_; pH 7.0) and added to the culture medium. Formed precipitates were carefully removed after 1 h. 600 μl of Lonza PNGM culture medium was added. For immunochemistry neurons were fixed for 7-10 min with 4% formaldehyde/4% sucrose in PBS at room temperature. After fixation, cells were washed three times in PBS and incubated for 1 h at room temperature with primary antibodies diluted in goat serum dilution buffer (GDB) (10% DS, 0,23% Triton X-100, in PBS). Neurons were then washed three times in PBS (5 min each), following incubation with Cy-conjugated secondary antibodies in GDB buffer for 1 h at room temperature. After three additional washes in PBS for 30 min each, slides were mounted using Vectashield mounting medium (Thermo Fisher Scientific, Dreieich, Germany). Images were acquired using a Nikon microscope equipped with the following components: Spinning Disk (Yokogawa) (Visitron Systems, Puchheim, Germany), solid state lasers (488, 561, 647 and 405), objectives (60x and 100x), two EM-CCD cameras (Hamamatsu Photonics 512/1024, Herrsching am Ammersee, Germany).

### Yeast two-hybrid-system

To map interaction domains of KIF21B and myosin Va, the lacZ-reporter gene assay of the DupLex-A yeast-two-hybrid system (Origene, Rockville, MD) was used. KIF21B-prey constructs encoding different domains were used in combination with bait constructs, encoding myosin Va domains.

### HSP induction and surface staining

At DIV 18-20 cultures were treated for 48 h with 50µM Bicuculline or 1µm TTX (Bio-techne GmbH, Wiesbaden, Germany). For surface labeling a GluA2 antibody was used, as described above. Cells were fixed for 4 minutes in 4% PFA. After washing 3 times with 1X PBS the primary antibody was added and incubated for 3 h at room temperature. After washing, cells were permeabilized with 0.1% Triton for 10 minutes. Phalloidin and the secondary antibody were added for 1 h at room temperature. Coverslips were mounted with Polymount (Polysciences Europe GmbH, Eppelheim, Germany) and imaged with a Nikon spinning disc confocal microscope (Visitron Systems, Puchheim, Germany).

### COS7 cell culture

Cells were cultured in Dulbecco’s modified Eagle’s medium (DMEM), (Thermo Fisher Scientific, Dreieich, Germany), supplemented with 10% fetal calf serum (FCS), 100 μg/ml streptomycin, and 100 units/ml penicillin. Cells were cultured at 37°C in a humidified incubator with 5% CO_2_ and seeded at 60,000 cells per 22 mm coverslip prior to transfection with Lipofectamine® 2000 (Thermo Fisher Scientific, Dreieich, Germany). Cells were used for time-lapse imaging experiments.

### Time-lapse imaging

COS7 cells or hippocampal neurons were prepared as described above. Coverslips were placed in an Attofluor® Cell chamber for microscopy (Thermo Fisher, Dreieich, Germany). Images were acquired using a Nikon microscope equipped with the following components: Spinning Disk (Yokogawa) (Visitron Systems, Puchheim, Germany), solid state lasers (488, 561, 647 and 405), objectives (60x and 100x), two EM-CCD cameras (Hamamatsu Photonics 512/1024, Herrsching am Ammersee, Germany) containing optical image splitters for simultaneous dual image acquisition, and an incubation chamber for controlled cell culture environment (5% CO_2_ at 37°C). Images were captured at 1-3 second intervals for 50-200 seconds.

### Fluorescence recovery after photobleaching (FRAP)

Cultured hippocampal neurons from *Kif21b* ^+/+^ and *Kif21b*^*–/–*^ mice were transfected at DIV 12 with GFP-actin plasmids and imaged the following day. Only spines with distinct heads were selected for analysis. Image acquisition was performed using a Nikon spinning disc confocal microscope (Visitron, Puchheim, Germany) equipped with 60x objectives, 405/488 nm lasers, and an incubation chamber (5% CO2, 37°C). Each spine was imaged five times (1s per frame) using 488 nm excitation before photobleaching. On the sixth frame, photobleaching (total bleaching time of 1 s) was induced with ∼2.2 mW of laser power (405 nm laser). Imaging resumed immediately after bleaching and continued every second for 300 consecutive seconds. The recovery of the bleached fluorescence signal for each frame was normalized to background levels and pre-bleach signals. The recovery curves were fitted to an exponential equation to extract various measures, including the relative size of the stable and dynamic pools as well as the recovery half-time (t_1/2_).

### Preparation of synaptosomes

Mice from both genotypes were euthanized, and brains were immediately extracted. Hippocampi were isolated and stored in 10 volumes of sucrose buffer 1 (320mM sucrose, 1mM NaHCO_3_, 1mM MgCl_2_, 0.5mM CaCl_2_, 1µM PMSF) containing protease inhibitor cocktail (Roche, Mannheim, Germany). Tissues were homogenized for 10 strokes using a motor-driven 2 ml Potter-Elvehjem homogenizer fitted with a Teflon pestle. All procedures were conducted at 4°C with pre-cooled solutions. Homogenates were centrifuged at 1,000 x g for 10 min. Resulting supernatants were stored on ice while pellets were re-homogenized once more, and centrifuged at 700 x g for 10 min. Resulting supernatants were combined with the first supernatant and centrifuged at 13,800 x g for 10 min. Pellets were homogenized in 500 µl sucrose buffer 2 (320 mM sucrose, 1 mM NaHCO_3_). Homogenates were overlaid with 2 ml of 1.4 M sucrose, 1 mM NaHCO_3_, and 2 ml of 1.0 M sucrose, 1 mM NaHCO_3_, followed by gradient centrifugation at 82,500 x g for 1.5 h. Bands at the 1.4 M and 1 M interface were collected, diluted in 4 volumes sucrose buffer 2, and pelleted by centrifugation at 28,000 x g for 20 min. The resulting pellets containing synaptosomes were re-suspended in 100µl sucrose buffer 2. For the preparation of PSDs, an equal volume of 1% (v/v) Triton X- 100, 320 mM sucrose, 12 mM Tris-HCl pH 8.0 was added for 15 min with occasional mixing. Samples were centrifuged at 70,000 x g for 1 h. Resulting supernatant were kept as PSD supernatant. The resulting pellet containing postsynaptic densities were resuspended in 40 mM Tris-HCl pH 8.0. Sample protein concentrations were determined with a BCA-assay (Pierce Biotechnology, Rockford, IL). Equal amounts of protein samples (10 µg of PSD supernatant and 5 µg of PSD) were applied on 4-15% gradient sodium dodecyl sulfate-polyacrylamide (SDS) gels.

### Electrophysiology

#### Slice culture preparation

Blind to genotype, organotypic hippocampal slices were prepared from WT and KO P5 mice as described previously^58^. Briefly, dissected hippocampi were cut into 400 μm pieces with a tissue chopper and placed on a porous membrane (2 slices per membrane, Millicell CM, Millipore) in six-well plates. Cultures were maintained at 37°C, 5% CO_2_ in a medium containing (for 500 ml): 394 ml Minimal Essential Medium (Sigma M7278), 100 ml heat inactivated donor horse serum (H1138 Sigma), 1 mM L-glutamine (Gibco 25030-024), 0.01 mg ml^−1^ insulin (Sigma I6634), 1.45 ml 5 M NaCl (S5150 Sigma), 2 mM MgSO_4_ (Fluka 63126), 1.44 mM CaCl_2_ (Fluka 21114), 0.00125% ascorbic acid (Fluka 11140), 13 mM D-glucose (Fluka 49152). No antibiotics were added to the culture medium. The medium was partially exchanged (60-70%) twice weekly. 36-48 hours before the electrophysiology recordings, membranes were placed into 35 mm culture dishes with fresh medium or fresh medium containing 40 µM bicuculline methiodide (Alamone labs, Jerusalem, Israel). The dishes were coded so that the experimenters were blind to genotype and treatment during the recordings and analysis.

Hippocampal slice cultures were placed at DIV 23-24 in the recording chamber and superfused with a HEPES-buffered solution containing (in mM): NaCl (145 mM, Sigma; S5886-500G), HEPES (10 mM, Sigma; H4034-100G), D-glucose (25 mM, Sigma; G7528-250G), KCl (2.5 mM, Fluka; 60121-1L), MgCl_2_ (1 mM, Fluka; 63020-1L), CaCl_2_ (2 mM, Honeywell; 21114-1L), pH 7.4, 318 mOsm kg^-1^. Patch pipettes with a tip resistance of 3 to 4 MΩ were filled with (in mM): K-gluconate (135 mM, Sigma; G4500-100G), EGTA (0.2 mM, Sigma-Aldrich; E0396-10G), HEPES (10 mM, Sigma; H4034-100G), MgCl_2_ (4 mM, Fluka; 63020-1L), Na_2_-ATP (4 mM, Aldrich; A26209-1G), Na-GTP (0.4 mM, Sigma; G8877-100MG), Na_2_-phosphocreatine (10 mM, Sigma; P7936-1G), ascorbate (3 mM, Sigma; A5960-100G), pH 7.2, 295 mOsm kg^-1^. TTX 1 µM, CPPene 1 µM, and picrotoxin 100 µM were added to the extracellular solution to isolate AMPA receptor-mediated miniature excitatory postsynaptic currents. Experiments were performed at 33°C ± 1 °C. Whole-cell patch-clamp recordings from CA3 pyramidal neurons were performed either using an Axopatch 200B (Axon Instruments, Inc.) amplifier or a Multiclamp 700B amplifier (Molecular Devices), both under the control of Ephus software written in Matlab (Suter et al., 2010). CA3 neurons were patched and held at −65 mV in the whole-cell voltage-clamp configuration (no liquid junction potential correction, LJP = -14.5 mV). Miniature EPSCs were recorded starting 5 min after break-in for 5 minutes. Series resistance, membrane resistance, and capacitance were calculated from −5 mV voltage steps (200 ms, every 100 s) from the holding potential of −65 mV. Series resistance was less than 20 MΩ, and recordings were discontinued if resistance changed more than 30%. The analog signals were filtered at 2 kHz and digitized at 10 kHz. *Analysis*: recordings were imported into Clampfit 10, high-pass filtered at 1 Hz, and events were detected using threshold detection (−12 pA). Statistical analysis was performed using GraphPad Prism.

### Diolistic (Dil) dye labeling in hippocampal slices

Mice were perfused with 4% PFA and 0.1% Glutaraldehyde in PBS. Hippocampi were sectioned into 300 μm slices, which were maintained in fixative until ready for use. *Preparation of bullets:* Diolistic neuronal staining was performed with 1.6 μM DiI-coated gold bullets were prepared by dissolving 13.5 mg of lipophilic Dil dye in 450 μl of methylene chloride (final concentration = 3 mg/100 μl). 100 μl DiI solution was dropped on 1 g of gold particles that were spread on a glass slide. After the methylene chloride had evaporated, the gold particles were scraped and diced with a razor blade, transferred to a tube containing 200μl water, and sonicated in a water bath for 10-30 min at room temperature. Meanwhile, TEZFEL tubing was coated with 10 mg/ml polyvinylpyrolidone (PVP) solution. Beads were added to the tubing via a syringe and rotated until all liquid had evaporated. The tubing was then cut into 13 mm-long sections and placed into a vial until use. *Shooting:* Dil-labeled bullets were shot through a 3.0 μm mesh 1 using a Helios Gene Gun system (Biorad, München, Germany) at 130 psi helium gas pressure.

### Image processing, spine head size

After Diolistic dye labeling, non-overlapping labeled neurons from the CA1-hippocampal region were imaged with a Nikon spinning disc confocal microscope (Visitron, Puchheim, Germany) using a 20x objective. Z-stack images were captured every 2 μm for subsequent three-dimensional reconstruction of the entire dendritic tree. In order to analyze spines, stacks of images with 1 μm step size were captured with a 60x objective. Secondary branches from basal and apical dendrites were selected as regions of interest from the original image stack using Imaris software (Bitplane, AG, Zurich, Switzerland). Using the filament tracer plugin, dendritic length, spine number and geometry of the spines were assessed. Values obtained for dendritic length, spine length, spine minimum diameter and spine terminal point diameter were used for further analyses. Spines with necks longer than 0.5 μm and head size of at least 0.13 μm^2^ were classified as mushroom spines. Only these spines were used for quantification of spine head size.

### Statistics

Statistical analyses were performed using SigmaPlot 13.0 (Systat Software Inc., Düsseldorf, Germany) and GraphPad Prism (GraphPad Inc., Bangalore, India).

## Author Contributions

KVG, MM and MK designed the study. KVG, ET, CEG, MS, DS performed experiments. All authors analyzed data. KVG and MK wrote the manuscript with help of MM. All authors read and commented on the manuscript.

## Acknowledgements

Supported by: Deutsche Forschungsgemeinschaft (DFG) grant KN556/11-2 (FOR 2419); KN556/16-1 and the Landesforschungsförderung Hamburg grant LFF-FV76 to M.K.

## Supplemental Information

**Figure S1.**
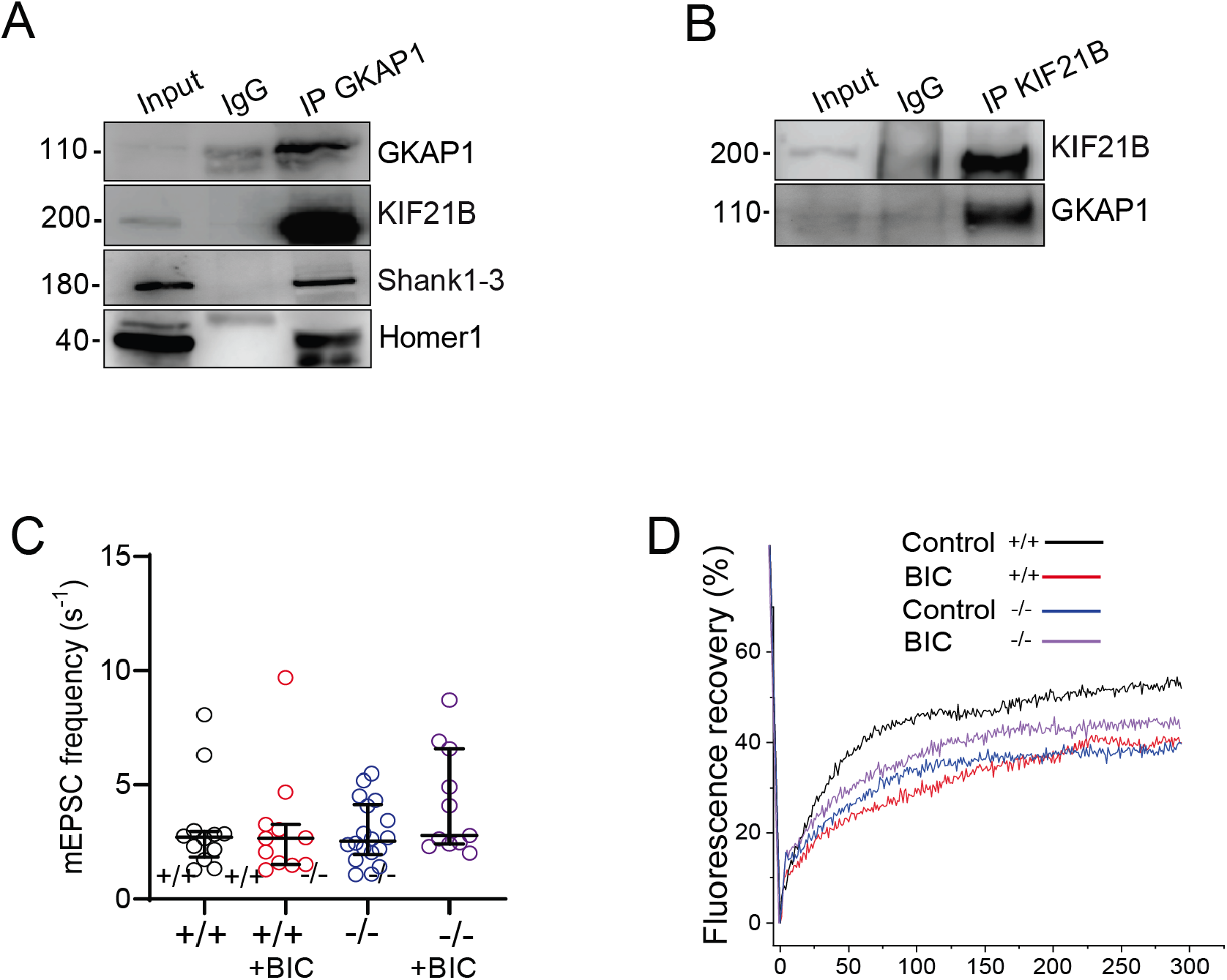
Supplemental Figure related to Figures 4 and 5. (A) Co-IP with GKAP-specific antibodies using adult mouse brain lysate. Detection of GKAP1, KIF21B, Shank 1-3(positive control), Homer-1 (positive control). N=3. (B) Co-IP with KIF21B-specific antibodies using adult mouse brain lysate. Detection of GKAP1, N=3. (C) Quantification of mEPSC frequency (n=11, 12, 18, 11) ANOVA, Post-hoc Tukey. p=0,4672 (D) FRAP analysis of GFP-actin recovery in spines of KIF21B^+/+^ or KIF21B^−/−^ neurons. Merged curves of Figure 5H and 5I.

**Movie S1.**
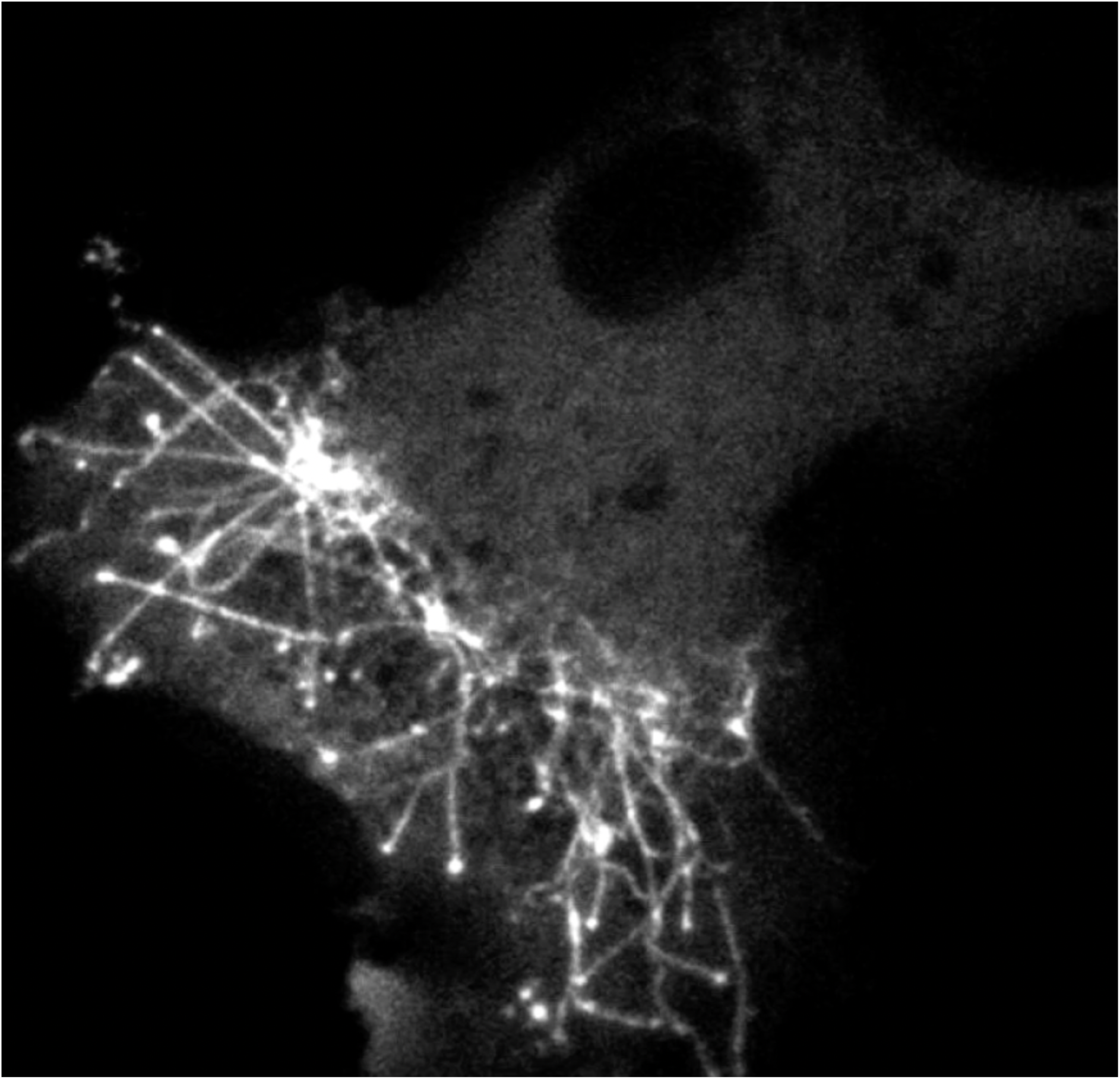
Still image of supplemental movie S1. Mobility of KIF2B-GFP particles in COS7 cells. Image acquisition intervals: 2 s over 3 min. The movie plays at 15 frames per second.

**Movie S2.**
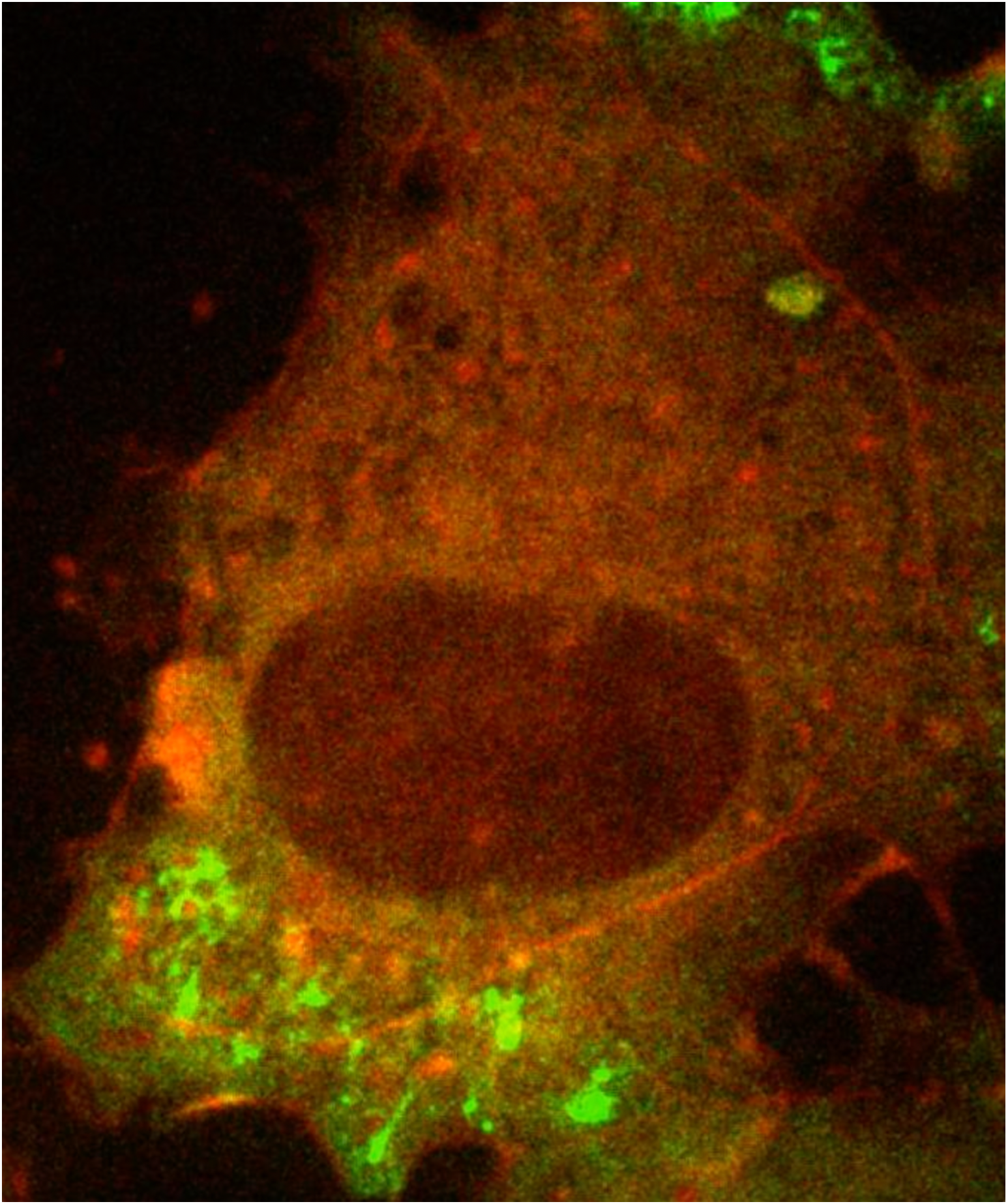
Still image of supplemental movie S2. Mobility of KIF2B-GFP and mRFP-actin in COS7 cells directly after cytochalasin D (CytoD) application. Image acquisition intervals: 2 s over 3 min. The movie plays at 15 frames per second.

